# Feeling the Music: Preceding Vibroacoustic Stimulation Modulates Oscillatory Brain Dynamics During Music Listening

**DOI:** 10.64898/2026.08.28.747481

**Authors:** Nandhini Natarajan, Iballa Burunat-Perez, Esa Ala-Ruona, Jan Kujala, Tiina Parviainen

**Author notes:** Data Availability Statement: Data available on request due to privacy/ethical restriction. Ethics Statement: This study received ethical approval from the Human Sciences Ethics Committee at the University of Jyväskylä on 28.08.23 (reference number: 1112/13.00.04.00/2023). Informed consent was obtained from participants electronically using the RedCap software and participants were made aware that they could withdraw at any time. Funding: This research was funded by the Research Council of Finland (grant number: 346210). The research is part of the Centre of Excellence in Music, Mind, Body and Brain. Conflict of Interest Disclosure: The authors have no conflicts of interest to disclose.

## Abstract

**Background:** Although typically considered an auditory experience, music listening engages multiple sensory systems, including somatosensory and motor pathways, making it an inherently multisensory phenomenon. However, research has predominantly examined the influence of music on other sensory systems, while the reciprocal question of how the existing state of a sensory system modulates the music listening experience, has received considerably less attention. To address this gap, we examined neural activity during music listening in two somatosensory states: one preceded by vibroacoustic stimulation (VAS) and one preceded by rest alone.

**Methods:** Forty participants completed two MEG sessions in a within-subject crossover design. In one session, they received 20 minutes of 40 Hz VAS before listening to 10 minutes of self-selected relaxing music (VAS_ML); in the other, they lay on the same mattress without stimulation (NoVAS_ML). Oscillatory and aperiodic activity were estimated using DICS beamforming and FOOOF decomposition for the whole music period and for early and late listening segments.

**Results:** Across the full listening period, the VAS condition was associated with reduced alpha power in the posterior temporal lobe and increased low-gamma power in the medial somatosensory and motor cortices compared to the NoVAS condition, suggesting enhanced cortical excitability and stronger auditory-motor engagement. Over time, both music listening conditions showed increases in alpha and beta power, consistent with habituation to the musical stimulus, though the spatial distribution differed qualitatively: changes were widespread across temporal and occipital regions in the NoVAS condition but remained localized to temporal areas after VAS. Additionally, VAS uniquely increased temporal-lobe theta power over time, whereas the NoVAS condition showed a decrease in the aperiodic exponent. Subjectively, participants reported stronger emotional intensity during music listening after VAS.

**Conclusion:** These findings suggest that preceding VAS induces a more engaged neural state and qualitatively alters the temporal dynamics of music processing.

## Introduction

Music engages various functional networks and regions in the brain. Not only does it (predictably) engage the auditory network (Han et al., 2025), it also engages the somatomotor, frontoparietal, dorsal attentional, reward and limbic networks (Alluri et al., 2017; Burunat et al., 2017; Chan & Han, 2022; Faber et al., 2023; Singer et al., 2016). Results from these studies indicate that music perception is not purely auditory, it also involves broader multisensory, cognitive and emotional networks. Music perception is often studied from the perspective of how the auditory external input of music influences our mental or motor states, including mental imagery (Jerling & Heyns, 2020; Taruffi et al., 2023), autobiographical memories (Belfi et al., 2015; Jakubowski & Ghosh, 2021) and urge to move (Duman et al., 2024; Janata et al., 2012). However, music listening (ML) experiences can also be approached from the ’opposite direction’, namely how music perception is influenced by our inherent body state. Our inherent body state is constantly modulated by various external stimuli. The somatosensory system, which continuously conveys information about external stimuli such as touch, pain, temperature etc is one such channel through which the internal bodily state can be modulated externally, for instance, through mechanical vibration applied to the body surface. ML experiences in concerts and clubs are often accompanied by low frequency sound vibrations, which form an integral part of the listening experience. Although previous studies have reported their effects on music perception, the neural correlates underlying these effects have not been extensively studied.

The literature on somatosensory stimulation generally uses two related approaches. Vibrotactile stimulation refers to mechanical vibrations delivered through contact surfaces such as platforms, chairs, or handheld devices, and is most commonly studied in perceptual and concert contexts. Vibroacoustic stimulation (VAS), by contrast, delivers low-frequency sound waves through a specialized mattress or therapeutic chair and is more prevalent in clinical and therapeutic settings (Leandertz et al., 2025; Punkanen & Ala-Ruona, 2012). The devices used across studies vary considerably, and this heterogeneity should be noted when comparing findings. Nevertheless, because the participants’ perceptual experience across these modalities is broadly similar, findings from vibrotactile studies are considered relevant to the current study. A study by Hove et al. (2020) found that listening to music along with vibrotactile stimulation leads to more forceful tapping to the music, more spontaneous body movement and higher ratings of enjoyment and groove. Another study by Siedenburg et al. (2024) that administered vibrotactile stimulation through surfaces of a chair along with music found that the stimulation enhanced the latent dimension of musical engagement which included groove, arousal and being a part of music. This mechanism allows creating more intense ML experiences, for example, in live or VR concerts. Cameron et al. (2022) found that low frequency sound processed via vibrotactile pathways increased dancing in live concerts. In the context of VR concerts, Venkatesan & Wang (2023) found that when participants listened to a concert with haptic feedback from a wristband, their sense of empathy, parasocial bond and loyalty towards the artist increased and feelings of loneliness decreased. This indicates that the somatosensory element of ML may also influence the emotional elements of the ML experience. Beyond its role as an accompaniment to music, tactile stimulation in the form of VAS has also been studied as a standalone intervention, where its effects on the body and mind can be examined independently of the music itself. This approach allows the specific contribution of somatosensory stimulation to the ML experience to be isolated. VAS finds applications in music therapy, where it is used as a grounding or relaxation technique and to enhance the effect of music (Fooks & Niebuhr, 2024; Leandertz et al., 2025; Punkanen & Ala-Ruona, 2012; Wheeler, n.d.). VAS has also been suggested to activate similar brain regions in participants who are hard of hearing as music does in participants with normal hearing (Lucía et al., 2020) and has also been found to enhance the activity of emotion and attention associated circuits in the brain (López et al., 2023). However, how preceding VAS alters oscillatory activity during naturalistic music listening in healthy individuals with normal hearing and how the effect evolves over time remains unknown. Although there is insufficient direct neuroscientific research on VAS and music, a study by Zabrecky et al. (2020) on participants with insomnia who received VAS found that VAS alters functional connectivity in various regions of the brain, including the prefrontal cortex, nucleus accumbens and sensorimotor areas. Furthermore, a study of 40 Hz VAS in mice found that VAS increases activity in the primary somatosensory and motor cortices (Suk et al., 2023). In this study, a vibroacoustic mattress will be used to administer the stimulation. The working hypothesis on how VAS influences music listening in the current study is that it modulates the somatosensory neural systems that are known to be involved in the processing of music (Alluri et al., 2017; Burunat et al., 2017).

Music listening is a temporally evolving process, with neural responses changing throughout the course of listening. Various studies have found that neural entrainment, i.e synchronisation of neural oscillations to periodic inputs, occurs to acoustic features of music, especially rhythm (Alluri et al., 2017; Burunat et al., 2017; Doelling et al., 2019; Fujioka et al., 2012; Tierney & Kraus, 2015). This entrainment seems to be particularly evident in the beta band in the auditory and motor cortices (Fujioka et al., 2012; Tierney & Kraus, 2015). However, most studies utilize simple sequences of notes or short pieces of music. fMRI studies on naturalistic music listening have also found that specific networks in the brain are modulated by specific acoustic features (Alluri et al., 2012; Burunat et al., 2017). Indeed, acoustic features and music listening are known to modulate neural activity with time, however, ML in the brain has not been explored from the perspective of higher level experiential states which may also vary along the same musical piece. Moreover, it is not known whether facilitating attention to the body via somatosensory stimulation by VAS modulates the brain state induced by music experience. To address this gap, we compared oscillatory activity during the first and last 3 minutes of a 10-minute ML experience following a vibroacoustic stimulation (VAS) or no stimulation (NoVAS) condition.

The neural power spectra have two distinct components: a broadband aperiodic signal which follows a 1/f distribution, and discrete oscillatory peaks arising at specific frequencies. The focus in past literature has been on periodic/oscillatory activity and aperiodic activity has mostly been ignored. However, recent studies report that aperiodic or broadband neural activity is physiologically meaningful and is modulated by factors such as task demands (Yan et al., 2024), age (Hill et al., 2022), diseases (Donoghue, 2025) and cognitive states (Frelih et al., 2025). Furthermore, it has also been suggested that some findings of oscillatory activity may not represent genuine oscillatory processes, but that apparent changes in oscillatory activity may instead reflect changes in aperiodic activity (Donoghue et al., 2020). To address this limitation, methods such as FOOOF (Donoghue et al., 2020) and IRASA (Wen & Liu, 2016) have been developed to characterise the periodic and aperiodic components more accurately. FOOOF (Fitting Oscillations and One Over F) (Donoghue et al., 2020) decomposes the neural spectra by first fitting and removing the aperiodic activity which is parameterized by its offset and exponent (slope of the spectra). It then models the residual peaks as Gaussians characterised by center frequency, power, and bandwidth. This enables us to determine if the recorded signal reflects an actual oscillatory peak or just elevated aperiodic activity, and allows the independent quantification of both.

Changes in oscillatory dynamics have consistently been found to reflect changes in physiological and cognitive states (Duda et al., 2024; Greene et al., 2023; Kang et al., 2011; Wu et al., 2026). In this study, we use changes in periodic and aperiodic activity as markers of shifts in brain state induced by VAS (VAS/NoVAS conditions) and listening phase (early vs late segments of listening to the musical piece). We conducted a within-subject magnetoencephalography (MEG) study with 40 participants across two experimental sessions. In the stimulation condition, participants received 20 minutes of 40 Hz VAS via a specially designed mattress, followed by resting-state recording, 10 minutes of self-selected ML, and a second resting-state recording. In the control condition, participants lay on the same mattress for the same duration without the stimulation, with all other procedures remaining the same.

We hypothesize that VAS will modulate brain state as reflected by oscillatory and aperiodic activity in the somatosensory and auditory regions and that these changes will evolve differently over time in the VAS and NoVAS conditions. This study provides insight into how sensory stimulation affects the temporal dynamics of music processing in the brain.

## Methods

### Participants

Forty healthy, right-handed participants (Age: 18-65 years; M= 33.08, 29 Females) with no history of neurological disorders or hearing loss were recruited through social media, word of mouth, and printed advertisements in Jyväskylä, Finland. All participants provided informed consent electronically prior to participation. The study was conducted in accordance with the ethical guidelines of the Finnish National Board on Research Integrity (TENK) and was approved by the Human Sciences Ethics Committee at the University of Jyväskylä.

### Experimental Design

The study comprised four sessions in total. The first two were accustomization sessions, conducted within one week of each other, during which participants visited the laboratory to experience VAS for 20 minutes. This was done to ensure that the participants are used to the feeling of the stimulation and the responses to it during the experiment are not to the novelty of the stimulation but the experience itself. VAS involved lying on a mattress containing low frequency transducers (Multivib mattress for clinic: https://multivib.com/en/home/) and receiving slow pulsatile stimulation at 40 Hz at an individually comfortable amplitude. This frequency was chosen as previous literature suggests that 40 Hz stimulation is relaxing and doesn’t interfere with other bodily rhythms like heart beat and respiration (Punkanen & Ala-Ruona, 2012). Self-reported measures of relaxation, arousal, and mood were collected on visual analogue scales before and after each accustomization session, and subjective reflection of thoughts, images, memories, emotions, and bodily sensations experienced during VAS and ML were recorded, though these are not reported here.

The final two sessions consisted of MEG recordings, separated by at least one week and beginning no sooner than one week after the last accustomization session. Each MEG session began with either a 20-minute period of VAS (VAS condition) or an equivalent period of lying on the vibroacoustic mattress without stimulation (NoVAS condition), with the order counterbalanced across participants. The MEG recording that followed comprised four blocks: an initial resting-state measurement (8 minutes eyes closed, 4 minutes eyes open), a 10-minute music listening task during which participants listened to self-selected relaxing music, a second resting-state measurement (8 minutes eyes closed, 4 minutes eyes open), and finally an interoception task. This study focuses on the differences in the neural activity during the ML task with and without stimulation. The resting state and interoception data are not reported here.

Behavioral measures of relaxation, arousal, and mood were collected at three time points: prior to the VAS or NoVAS block (T1), immediately following it (T2), and after the ML task (T3). Although the experiential measures at last time-point (T3) were collected after the MEG recording had concluded, participants were asked to retrospectively report their affective state during the ML task; this approach was adopted due to time constraints within the MEG recording protocol. To compare the ML experience across the VAS and NoVAS conditions, participants rated the music on four dimensions: valence (whether it affected them positively or negatively), arousal (whether it increased or decreased their energy), the intensity of the emotions they felt during listening and familiarity to the music. These ratings were collected after the MEG recording.

### Music Stimuli

Participants chose two pieces of music they considered relaxing, each at least five minutes in duration. One of these was selected as the subject-specific music stimulus based on its availability and cost on the Apple Music store. The selected track was then looped twice using the BandLab software (BandLab Technologies, Singapore, https://www.bandlab.com) to produce a stimulus file of sufficient length, which was edited so that the audio faded out after ten minutes.The same track was used in both, the VAS and NoVAS, conditions.

### MEG Data Recording

Neuromagnetic activity was recorded using a whole-head 306-channel MEG system comprising 102 magnetometers and 204 planar gradiometers (TRIUX, MEGIN Oy, Helsinki, Finland), housed within a magnetically shielded room at the Centre for Interdisciplinary Brain Research, University of Jyväskylä. Five head-position indicator coils attached to the scalp, with three positioned on the forehead and one behind each ear, were used to monitor the head position. Electrooculography (EOG) using two electrodes, placed above the right outer canthus and below the left outer canthus respectively, was used to monitor ocular activity. Electrocardiography (ECG) using three electrodes positioned below the right clavicle, below the left clavicle, and on the right clavicle as ground was used to monitor cardiac activity. To facilitate co-registration with a standard MRI template, three anatomical fiducial points (the nasion and the left and right preauricular points) along with approximately 150 additional scalp surface points were digitized to define the head coordinate system. MEG data were sampled at 1000 Hz.

### Preprocessing

The data were first processed with MaxFilter (version 2.2, MEGIN Oy, Helsinki, Finland) to identify bad channels, suppress external artefacts, and realign recordings to each participant’s median head position. The cleaned signals were then band-pass filtered between 1-40 Hz and downsampled to 200 Hz. Independent component analysis (30 components) was applied using MNE (version 1.6.1; Gramfort et al., 2013) and components reflecting eye blinks and cardiac activity were removed following visual inspection of their field patterns and time-courses.

### Source Analysis

In the MEG data analysis, we estimated the power spectra for six different samples from the music listening condition: the whole 10 minutes (whole music piece) as well as the first and last three minutes of the data for both the VAS and NoVAS condition (early and late segments).

Dynamic Imaging of Coherent Sources (DICS; Gross et al., 2001) was used to map periodic and aperiodic neural activity during music listening in both stimulation conditions. Cross-spectral density (CSD) matrices for 1–40 Hz were first computed using Welch’s method (2048-sample windows, 50% overlap, Hanning taper). These CSDs served as input to a DICS beamformer to estimate the cortical distribution of oscillatory and broadband activity, implemented with custom MATLAB scripts (R2020b, The MathWorks Inc.).

DICS was performed at the parcel level using a template brain (fsaverage-5.1.0) and a single-compartment realistic boundary element model (Gramfort et al., 2014). Cortical parcels were defined using a modified Destrieux atlas (Destrieux et al., 2010) comprising 69 parcels in the left and 68 parcels in the right hemisphere, adjusted for approximately uniform size (Ala-Salomäki et al., 2021).

A frequency-domain beamformer with common spatial filters was constructed from CSDs pooled across all conditions. Here, a broadband beamformer was constructed in the 2-40 Hz range, and the obtained beamformer weights were averaged across the source points belonging to each parcel. These average weights were then applied to each frequency bin separately for each condition to obtain parcel-wise power spectra for the whole 10 (VAS_ML, NoVAS_ML), the first three (VAS_early_ML, NoVAS_early_ML) and the last three minutes (VAS_late_ML, NoVAS_late_ML) of the data.

Finally, the FOOOF algorithm (Donoghue et al., 2020) was used to decompose each parcel’s spectrum into aperiodic (exponent, offset) and periodic components using the default options (peak width: 0.5-12 Hz; absolute peak threshold: 0; relative peak threshold: 2; aperiodic mode: fixed). Periodic power was derived by subtracting the aperiodic fit from the full spectrum and averaging the residual power within canonical theta (4–7 Hz), alpha (8–13 Hz), low beta (13–20 Hz), high beta (20–30 Hz), and low gamma (30–40 Hz) bands.

### Statistical Analysis

Parcel-wise differences in periodic and aperiodic neural activity between conditions were assessed separately for the left and right hemispheres using cluster-based permutation tests. Prior to band-averaging, the aperiodic component of each parcel’s power spectrum was removed by subtracting the FOOOF-derived aperiodic fit, ensuring that subsequent analyses of oscillatory power reflected purely periodic activity. Mean periodic power was computed for five frequency bands: theta, alpha, low beta, high beta, and low gamma. Aperiodic parameters (exponent and offset) were analysed separately using the same pipeline.

At each parcel, a paired-samples t-test was used to compare the difference between conditions (VAS_ML - NoVAS_ML, VAS_late_ML - VAS_early_ML, NoVAS_late_ML - NoVAS_early_ML) across participants. Parcels were considered candidate cluster members if they exceeded a threshold of p < 0.05 and were of the same sign (positive and negative t-values were clustered separately). Weighted clustering was applied to group parcels into clusters using the Euclidean distance of 4 cm across parcel coordinates. The cluster statistic was defined as the sum of t-values across all parcels within a cluster.

Statistical significance was assessed via a sign-flipping permutation procedure (5,000 permutations). On each permutation, the condition difference scores of a randomly selected subset of participants were sign-flipped, effectively reassigning condition labels within the within-subjects design. The cluster-forming and clustering procedure was repeated for each permuted dataset, and the maximum positive and minimum negative cluster t-sum were recorded. A null distribution was constructed by pooling the maximum positive and absolute minimum negative cluster statistics across all permutations, leading to a distribution of 10,000 cluster-level t-scores. The observed cluster statistic was compared against the 95th percentile of this null distribution (cluster-level threshold: p < 0.05). To characterise the magnitude of significant cluster effects, mean power was extracted per participant across each significant cluster and averaged within each condition. Descriptive statistics (mean, SD) and effect sizes (Cohen’s d) were computed from these values. All analyses were performed separately for the left and right hemispheres using custom scripts in MATLAB (R2020b).

Custom Python scripts were used to perform statistical analyses of the music measures collected post ML. The normality of each variable was visually inspected using histograms and statistically tested with the Shapiro-Wilk test. The data was found to be normally distributed and paired t-tests were performed to assess the differences in valence, arousal, felt intensity of emotions and familiarity to the music.

## Results

### Whole Music Piece Comparison (VAS_ML - NoVAS_ML)

Cluster permutation test revealed that alpha power was lower in the VAS_ML condition compared to the NoVAS_ML condition in the posterior temporal lobe. Low gamma power was found to be higher in the medial somatosensory and motor cortex in the VAS_ML condition compared to the NoVAS_ML condition. (Fig 3, Table 1)

**Fig 1:**
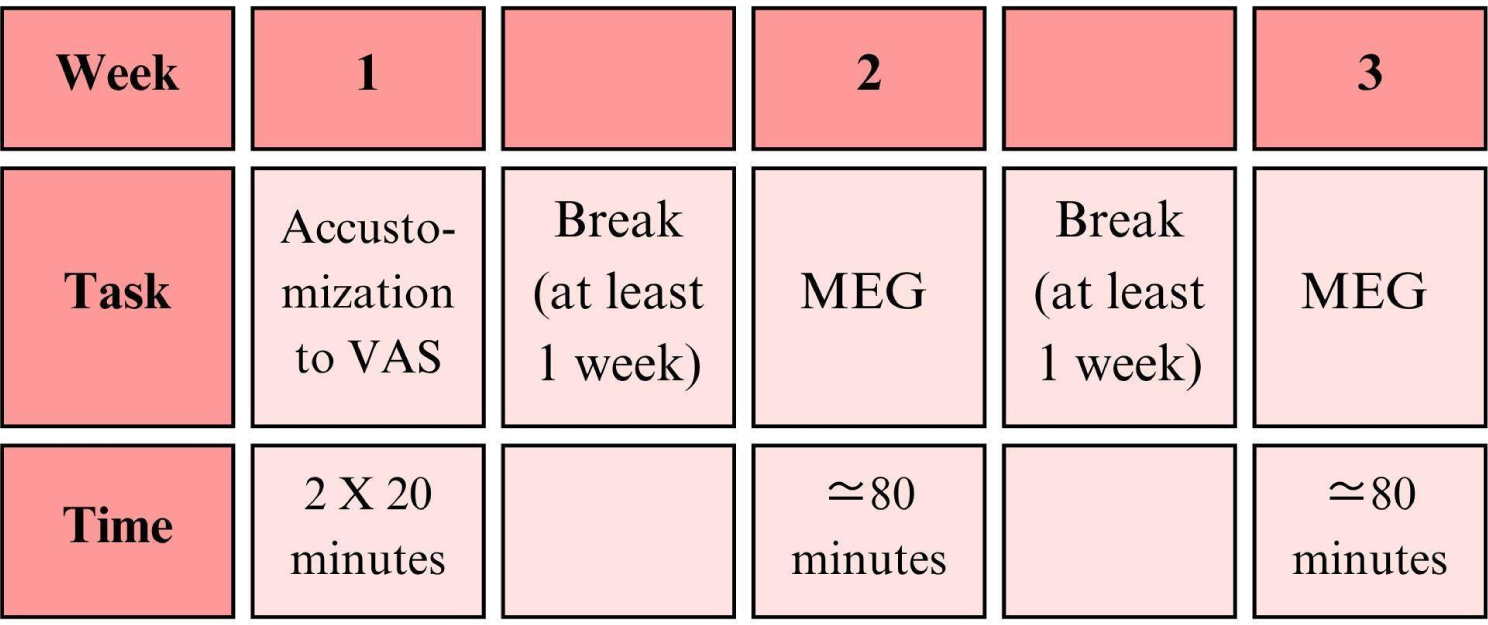
Overview of study.

**Fig 2:**
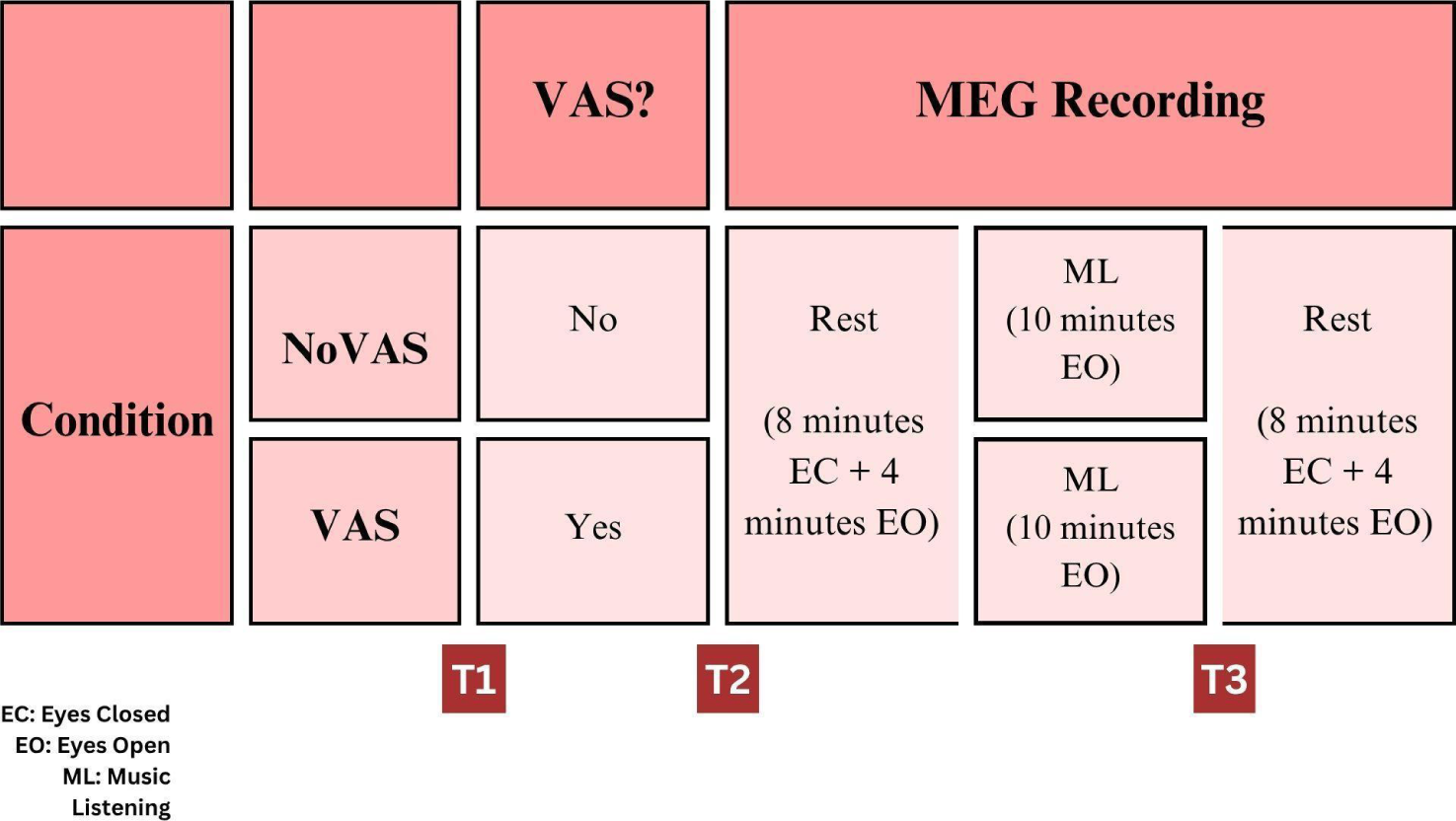
Overview of MEG session.

**Fig 3:**
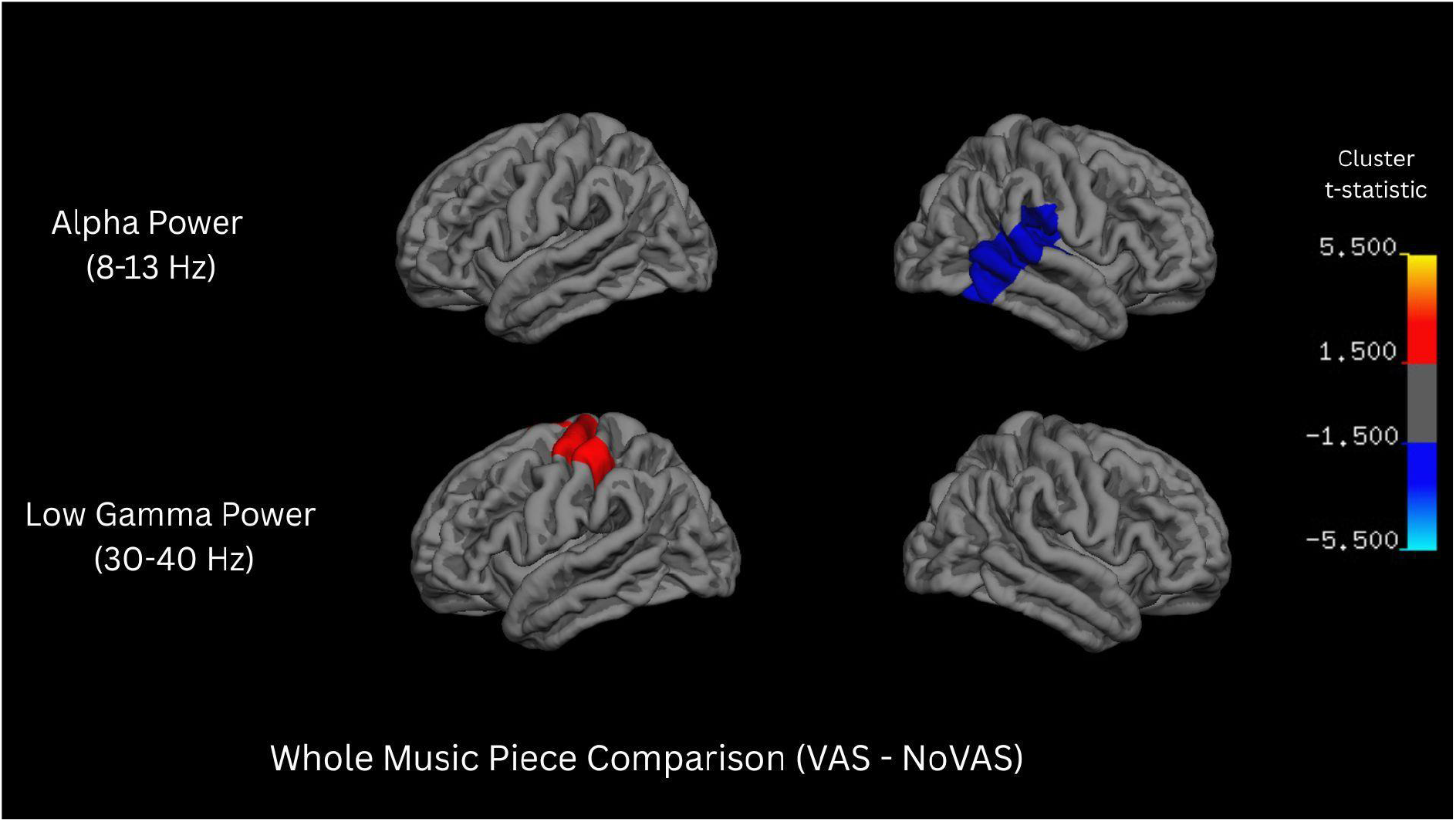
Oscillatory power differences between the VAS and NoVAS conditions during the whole listening session.

**Table 1:**
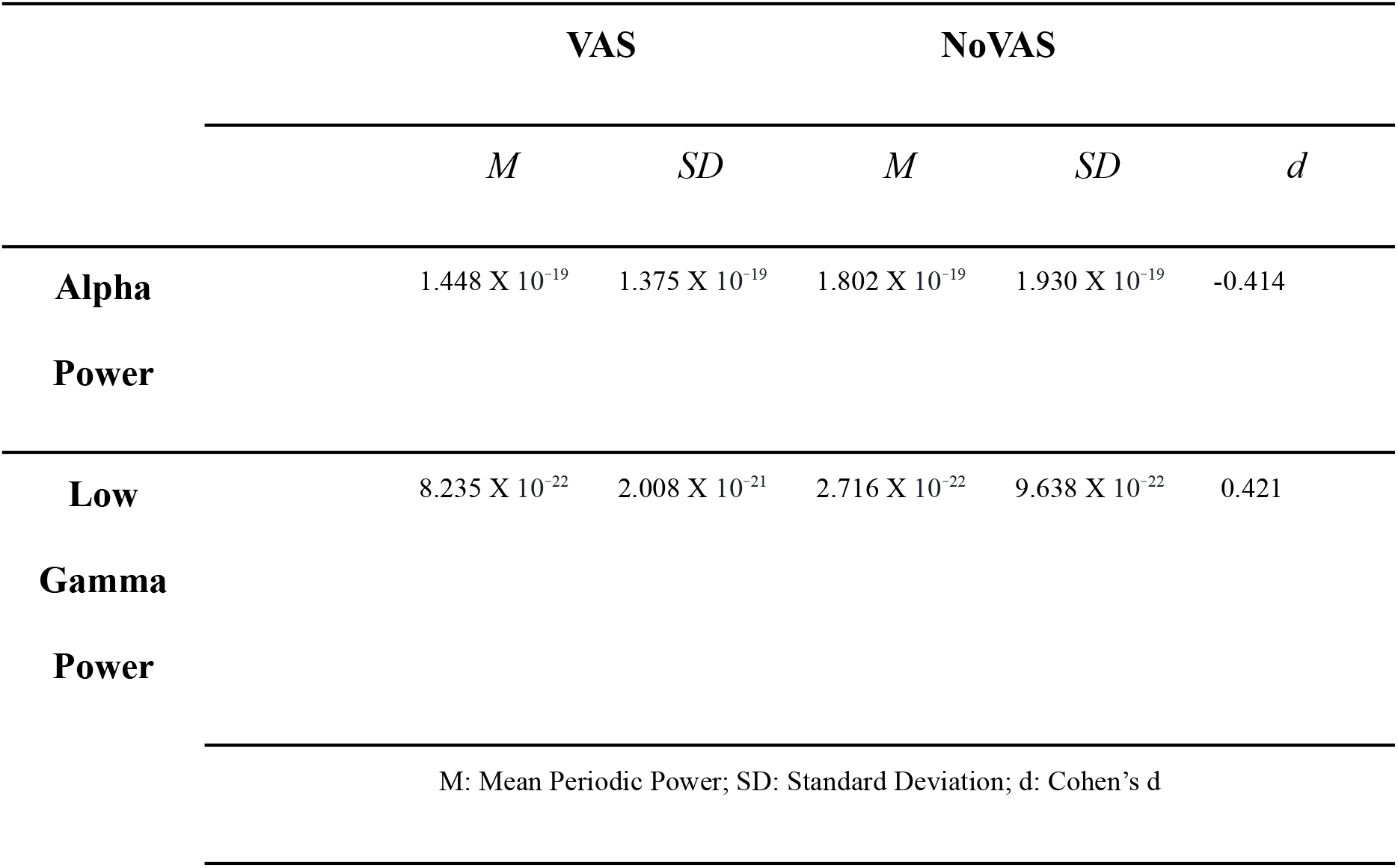
Cluster-level descriptive statistics for the VAS and NoVAS conditions.

### Segment Comparison (VAS_late_ML - VAS_early_ML and NoVAS_late_ML - NoVAS_early_ML)

In the NoVAS condition, cluster permutation test revealed that the exponent decreased over time in the left inferior frontal gyrus during ML. Alpha power was found to increase with time in the left middle and posterior temporal lobe and the right occipital lobe over time during ML. High beta power increased with time in the occipital lobe bilaterally but asymmetrically. (Fig 4; Table 2)

**Fig 4:**
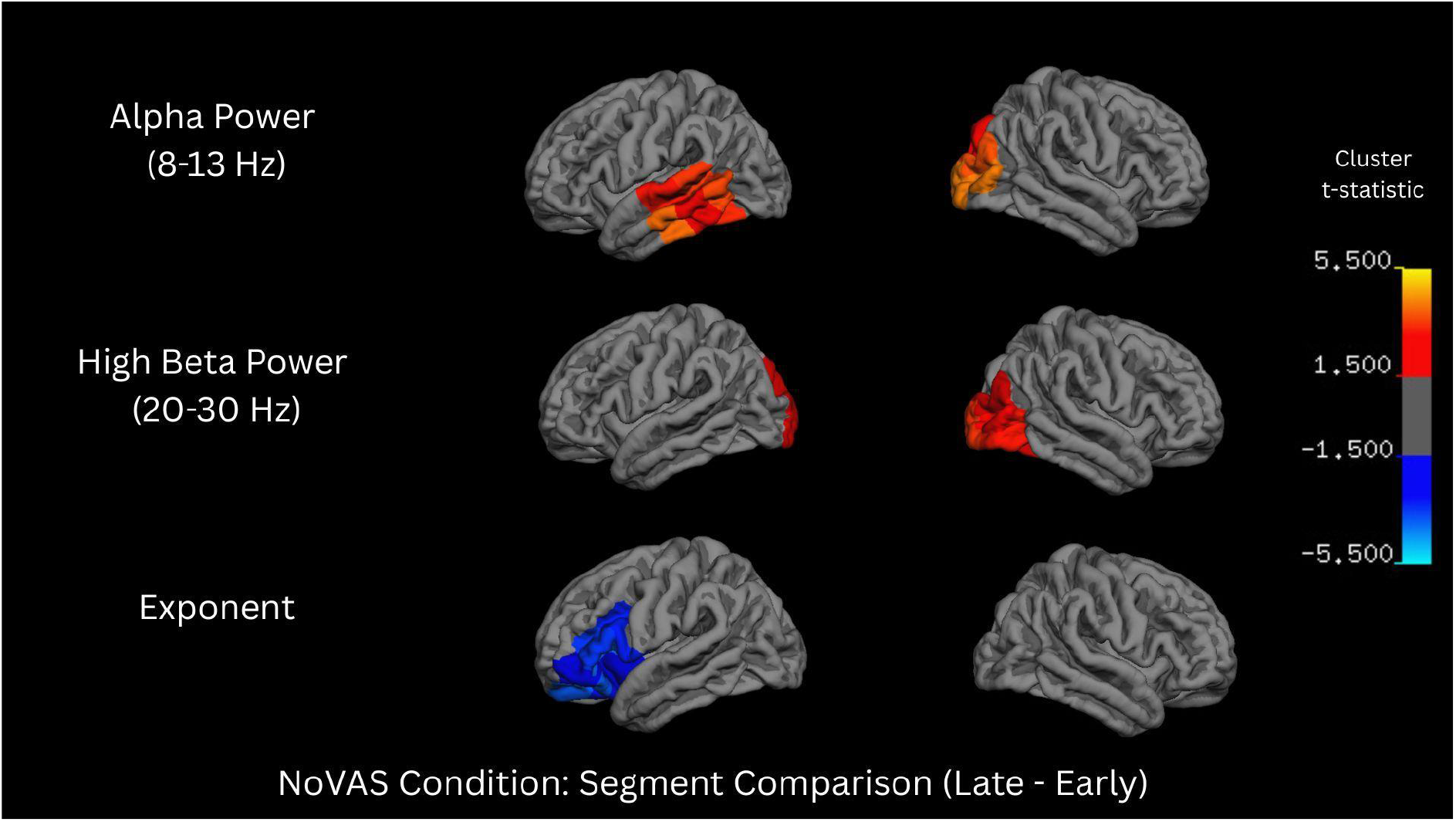
Oscillatory Power and Aperiodic exponent differences between the late and early segments of music listening in the NoVAS condition.

**Table 2:** Cluster-level descriptive statistics for the early and late segments of the NoVAS condition.

|  |  | Early Segment |  | Late Segment |  |  |
| --- | --- | --- | --- | --- | --- | --- |
|  |  | *M* | *SD* | *M* | *SD* | *d* |
| **Exponent** | Cluster 1 | 0.859 | 0.128 | 0.819 | 0.136 | -0.572 |
| **Alpha** | Cluster 1 (TL) | 1.296 X 10<sup>-19</sup> | 1.475 X 10<sup>-19</sup> | 1.609 X 10<sup>-19</sup> | 1.609 X 10<sup>-19</sup> | 0.537 |
| **Power** | Cluster 2<br>(OL) | 2.659 X 10<sup>-19</sup> | 3.237 X 10<sup>-19</sup> | 3.343 X 10<sup>-19</sup> | 3.985 X 10<sup>-19</sup> | 0.496 |
| **High Beta** | Cluster 1<br>(Left) | 3.012 X 10<sup>-20</sup> | 3.368 X 10<sup>-20</sup> | 3.540 X 10<sup>-20</sup> | 3.920 X 10<sup>-20</sup> | 0.447 |
| **Power** | Cluster 2<br>(Right) | 2.093 X 10<sup>-20</sup> | 1.621 X 10<sup>-20</sup> | 2.448 X 10<sup>-20</sup> | 2.072 X 10<sup>-20</sup> | 0.457 |
| M: Mean Periodic Power; SD: Standard Deviation; d: Cohen's d; TL: Temporal Lobe; OL: Occipital Lobe |  |  |  |  |  |  |

In the VAS condition, theta power was found to increase with time in the left posterior superior temporal gyrus and middle and inferior temporal gyrus. Alpha power increased over time in the left temporal pole and inferior temporal gyrus and part of the inferior occipital gyrus during music listening. Low beta power was found to increase over time in the left middle temporal lobe during music listening. High beta power also increases with time bilaterally in parts of the temporal lobe. (Fig 5; Table 3)

**Fig 5:**
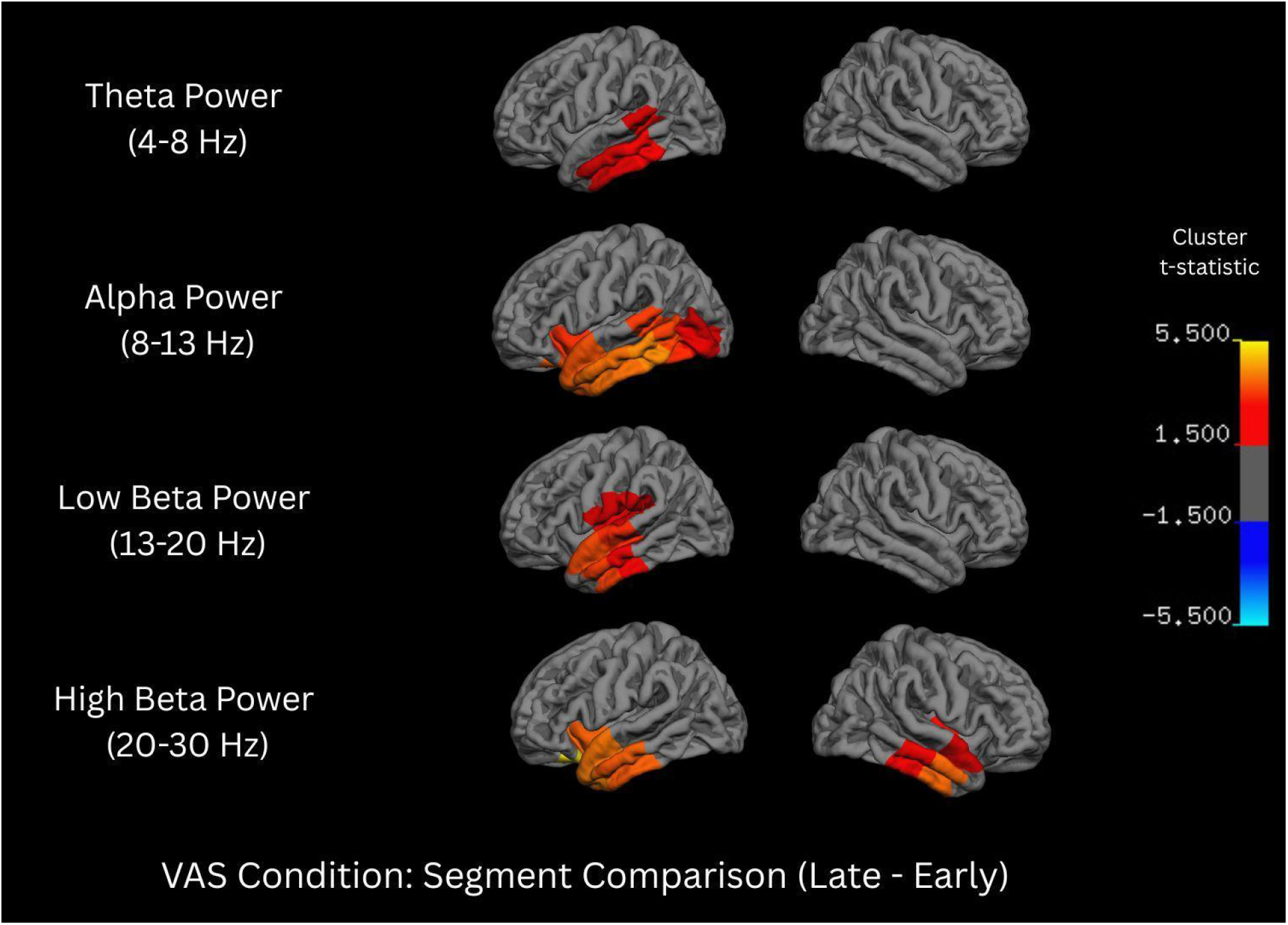
Oscillatory Power differences between the late and early segments of music listening in the VAS condition.

**Table 3:**
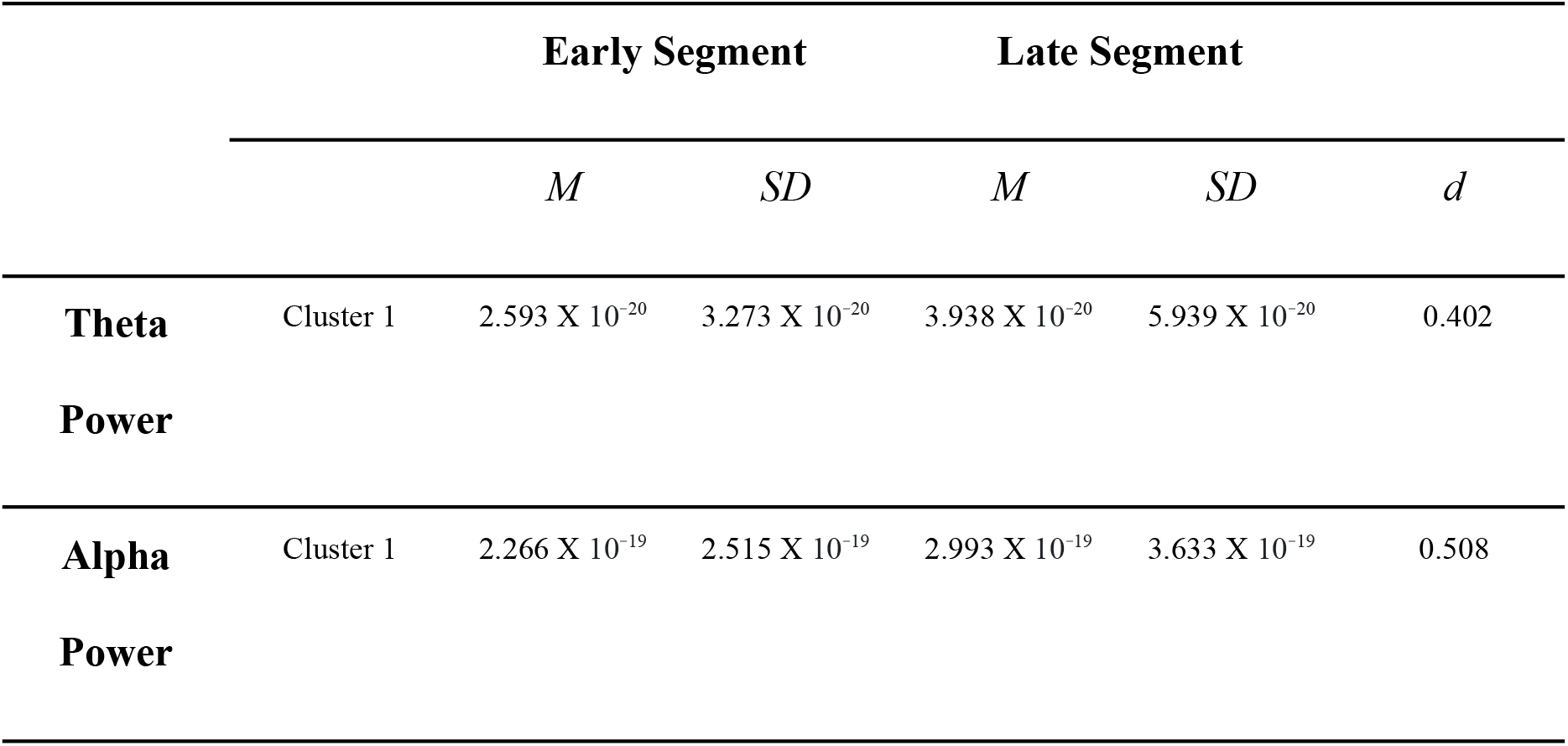
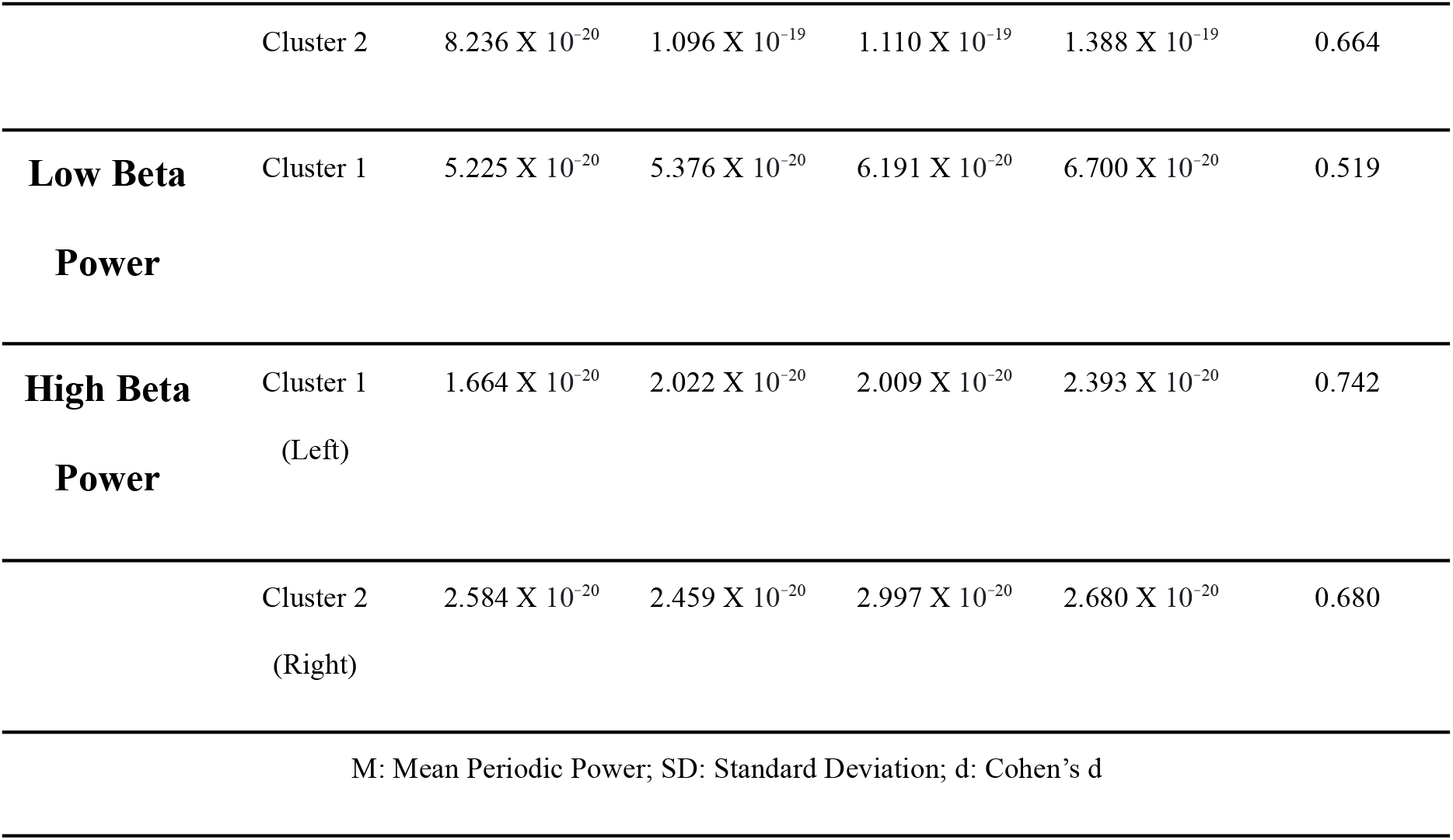
Cluster-level descriptive statistics for the early and late segments of the VAS condition.

### Music Measures

Paired t-tests revealed a significant difference in the intensity of felt emotions in the VAS_ML condition compared to the NoVAS_ML condition. Participants reported that they felt the emotions more strongly in the VAS_ML condition compared to the NoVAS_ML condition (t = 2.12, p< 0.05, d=0.337). No significant differences were found for valence, arousal or familiarity to the music.

## Discussion

This study examined differences in periodic and aperiodic activity during ML between two stimulation/somatosensory conditions, namely with and without preceding VAS (VAS_ML vs NoVAS_ML), and between early and late segments of ML (VAS/NoVAS_early_ML vs VAS/NoVAS_late_ML). Comparison across the whole music piece revealed lower alpha power in the PTL and higher low-gamma power in the medial primary somatosensory and motor cortices in the VAS_ML compared to the NoVAS_ML condition. Over time, alpha and beta activity increased in both the VAS and NoVAS conditions though the spatial distribution of the increase was different. In the NoVAS condition, alpha increase occurred both in the temporal and occipital regions whereas high beta increase occurred in the occipital regions. Whereas, the increase in alpha and beta power were localized to temporal regions in the VAS condition. In addition, in the NoVAS condition the aperiodic exponent decreased over time and in the VAS condition the theta power increased over time.

Two main effects were studied (VAS_ML - NoVAS_ML; Late - Early), each addressing a different question about how VAS modulates brain states induced by ML. The whole session comparison captures the mean oscillatory state while the segment comparison analysis captures dynamic trajectories within the session.

### Effect of VAS on the ML Experience (Whole Session Comparison)

As compared with the NoVAS condition, the alpha power in the posterior temporal lobe was lower during ML after VAS. According to the Gating by Inhibition theory (Jensen & Mazaheri, 2010; Klimesch et al., 2007), alpha activity increases in task irrelevant regions and enables gating of information to task relevant regions. A decrease in alpha in this region, which is thought to be involved in higher order sensory processing, could indicate that cortical engagement increases during ML after VAS relative to the NoVAS condition. In other words, music-induced alpha decrease is more pronounced in posterior temporal areas after VAS than after mere rest. Furthermore, we found an increase in low gamma power in the medial primary somatosensory and motor cortices which may reflect increased sensorimotor involvement during ML. In line with this interpretation, somato-motor network has been consistently found to be recruited during beat perception even in the absence of movement (Chen et al., 2008; Grahn & Brett, 2007). Taken together, we can speculate that the stimulation may induce a state where music can be more actively processed by reducing inhibition of auditory processing areas in the PTL and increasing activity in regions of the auditory-motor network.

### Common Oscillatory Changes in Conditions over Time (Segment Comparison)

Alpha and beta power both increased in the regions of the temporal lobe. Both of these frequency bands are associated with habituation and maintenance of somatosensory state. It would be interesting to interpret their increase over time in the present study as reflecting brain habituation to the music stimuli over time. Alpha power typically decreases on presentation of a novel stimulus (Amochaev et al., 1989). Similarly, beta oscillations have been associated with the maintenance of the somatosensory or cognitive state with increase in activity when a change is not expected (Engel & Fries, 2010). Additionally, beta band activity is associated with working memory and temporary storing of information before recall (Spitzer & Haegens, 2017). Our results are partially in line with Höller et al. (2012), who were examining common neural responses to self selected music and found that even though there were interindividual differences, there were some subjects who experienced synchronization in the higher alpha and beta power. Interpreting our findings in light of those earlier results, listening to music over time may induce habituation-like neural adaptation, and the cognitive engagement needed to process the music may decrease. These results and interpretation are also in line with Jäncke et al., 2015 who found an increase in oscillatory activity in alpha, beta and theta bands over the course of ML and between rest and ML but not during repeated presentation of the same stimulus. This could indicate that the increase in oscillatory activity in this case is due to habituation and not familiarity. In fact, familiarity is associated with a suppression and not increase of alpha and beta activity (Malekmohammadi et al., 2023).

### Oscillatory and Aperiodic Changes in the VAS and NoVAS Conditions Over Time (Segment Comparison)

Although the direction of oscillatory changes at alpha and beta bands was the same in the VAS and NoVAS conditions, the spatial distribution of the increase in oscillatory activity was qualitatively more widespread in the NoVAS than in the VAS condition. In the NoVAS condition, the increase was in both the temporal and occipital lobes, whereas in the VAS condition, the increase was localized to the temporal lobe.

We speculate that in the NoVAS condition, ML potentially induces a more idling-like state, similar to activity profiles of participants at rest (Berger, 1929; Brookes et al., 2011), whereas in the VAS condition, the brain maintains a more active and engaged state. Furthermore, we also found an increase in theta activity in the left middle and inferior temporal lobe. Increases in theta during cognitive tasks is generally indicative of better task performance (Tan et al., 2024). Theta activity is also associated with encoding of memories (Nyhus et al., 2019) and meditative states (Duda et al., 2024; Lagopoulos et al., 2009). Additionally, this finding points to the fact that the stimulation changes the qualitative nature of the neural correlates of the ML experience and not just the magnitude or spatial extent.

Analysis of aperiodic activity using FOOOF revealed that the aperiodic exponent decreased over time in the NoVAS condition, reflecting a progressive flattening of the 1/f slope during ML. This pattern may be consistent with a shift towards relatively greater excitation or reduced inhibition within cortical circuitry, although the excitation-inhibition interpretation remains indirect in MEG data. No such change was observed in the VAS condition. This could also mean that the brain is more likely to show broadband fluctuations in the NoVAS condition whereas VAS maintains more structured neural signalling. Critically, because periodic and aperiodic components were modeled independently, the oscillatory findings reported above cannot be attributed to broadband aperiodic changes, i.e the alpha, beta, and theta effects reflect genuine periodic activity.

Jointly, the whole session comparison revealed reduced alpha power in the PTL and increased low gamma power in the medial sensorimotor regions following VAS, indicating that VAS is associated with a more engaged and active brain state. From the segment comparison, we found an increase in alpha and beta activity in both conditions, suggesting that that could be a general response of the brain to music over time. The brain possibly predicted the incoming stimulus better and habituated to it. The differences in condition over time were the spatial extent of alpha and beta activity changes, theta increase in the VAS condition and exponent decrease in the NoVAS condition. In the VAS condition, the changes were concentrated in the temporal lobe, whereas in the NoVAS condition, oscillatory activity increased in the temporal and occipital lobes over time. Increase in alpha and beta activity in the occipital lobe is generally indicative of a rest-like idling state (Berger, 1929; Brookes et al., 2011). Considering this and the increase in theta in the VAS condition and decrease in exponent in the NoVAS condition, we suggest that the VAS lead to a state where the music was processed more actively and was possibly embodied more than without VAS. Interestingly, the subjective ratings also indicate that the emotions felt through the music were stronger in the VAS relative to the NoVAS condition.

These results provide valuable additions to the literature on VAS and ML. To our knowledge, this is the first study examining the neural correlates of ML in two sensory conditions. Although previous studies on the use of vibrotactile stimulation in VR concerts (Venkatesan & Wang, 2023) and the neural correlates of vibrotactile stimulation in people who are hard of hearing (Lucía et al., 2020) have also found that stimulation can be used to enhance the ML experience, this study supports those claims by studying healthy individuals in a crossover design with and without stimulation. Additionally, since we used self-selected relaxing music as the stimulus and the effect was averaged over 10 minutes in the whole session comparison and 3 minutes in the over time comparison, we can, with a degree of certainty, say that the effect was not music dependent. These findings are promising and may find applications in the novel design of devices that can aid the ML experience of deaf/hard hearing individuals, in music therapy and in creating more intense and immersive live ML experience for the public.

### Limitations and Future Directions

Although this study offers valuable insights into the effects of vibroacoustic stimulation and music listening, several limitations should be acknowledged. First, VAS was administered *before* music listening rather than concurrently. This choice allowed us to isolate the effect of the stimulation and avoid introducing artefacts into the MEG signal, as the VAS device was not MEG-compatible and could have generated noise during music listening. However, future studies should examine both, administering VAS before and during the ML to gain a causal understanding of the effect of VAS on the ML experience. Second, individual MRI scans were not available for all participants, which limits the anatomical precision of the reported spatial distributions. Third, there could have been individual differences in the experience of VAS or ML that could not be captured. These factors should be considered when interpreting the findings, and future studies would benefit from addressing these methodological constraints. Future studies should also examine connectivity differences between these conditions to further understand the dynamics of these differences.

## Conclusions and Implications

Preceding vibroacoustic stimulation altered the neural dynamics of subsequent music listening. It was associated with increased auditory cortical engagement and enhanced auditory-motor involvement throughout the listening session. It also increased habituation-related oscillatory dynamics within the auditory network and it progressively recruited a theta-based process that listening in the NoVAS condition did not engage. Together, these findings suggest that vibroacoustic stimulation did not merely change the magnitude of neural responses to music listening, but also altered it qualitatively– shifting the brain from passive listening towards a more engaged and embodied listening state.

## Acknowledgements

The authors would like to thank Joonas Hyttinen for support with data collection; Simo Monto and Viki-Veikko Elomaa for their consistent guidance and technical support; and all the participants for their time.

